# Boosting the signal: Expectation-driven gain modulation of preparatory spatial attention

**DOI:** 10.64898/2025.12.10.693414

**Authors:** Dirk van Moorselaar, Stefan Van der Stigchel

**Affiliations:** Faculty of Social and Behavioral Sciences, Experimental Psychology, Helmholtz Institute, Utrecht University, Utrecht, the Netherlands

## Abstract

The visual system can flexibly adjust attentional deployment to match task demands, but whether observers can proactively modulate the spatial scope of attention based on expectations about upcoming search difficulty remains unclear. According to the zoom lens model, attention can narrow or broaden its spatial extent, with narrower focus enhancing processing efficiency, a mechanism that would benefit target discrimination in crowded displays. We tested whether observers adjust attentional scope when expecting sparse versus dense search arrays by combining spatial cueing with block-wise manipulations of display density expectations. Participants performed a visual search task in which endogenous cues predicted target location, while blocks predominantly contained either sparse (1 target, 3 distractors) or dense (1 target, 7 distractors) displays. We applied inverted encoding models to broadband EEG data to reconstruct spatial channel tuning functions, enabling precise characterization of both the locus and breadth of attentional deployment. Behaviorally, expecting difficult searches selectively improved accuracy at cued locations without costs elsewhere. Consistent with this selective benefit, neural measurements revealed that expectancy enhanced the amplitude of spatially selective responses at the attended location but did not alter tuning width. These findings demonstrate that expectations about search difficulty modulate attention through gain-based signal enhancement rather than adjustments to spatial scope, revealing that preparatory attentional control operates via amplitude modulation within a stable spatial focus. This mechanism complements reactive attentional adjustments and provides an efficient means for the visual system to optimize processing under predictable task demands.

## Introduction

Whether navigating a crowded street or searching for a single item in an organized drawer, our visual system continuously adapts the scope of attention to match task demands. Early spatial cueing paradigms conceptualized attention as a spotlight with fixed dimensions that shifts across space (Posner, 1980; Posner et al., 1980). However, the influential Zoom Lens Model proposed a more flexible framework wherein attentional resources dynamically distribute across different spatial scales (Eriksen & St. James, 1986). This model posits that attention spreads with a gradient of processing quality decreasing from its center, and critically, an inverse relationship between attentional scope and perceptual enhancement: narrower focus yields greater processing efficiency at the attended location (Castiello & Umiltà, 1990). Empirical support demonstrates that narrowing attention both enhances behavioral performance (Eriksen & St. James, 1986) and modulates neural activity in early visual cortex (Heinze et al., 1990; Müller & Kleinschmidt, 2003), with tuning profiles in alpha-band oscillations tracking the breadth of attentional focus (Feldmann-Wüstefeld & Awh, 2020; van Moorselaar et al., 2025).

Beyond adjusting the spatial extent of attentional deployment in response to immediate perceptual input, the visual system can proactively configure attention based on learned expectations about upcoming stimulus displays. Research on preparatory attentional control has demonstrated that observers flexibly adjust attentional settings according to task demands (Belopolsky et al., 2007) and can adjust preparatory attentional tuning in response to spatial probability learning (van Moorselaar & Slagter, 2019). Critically, such preparatory adjustments occur before stimulus onset, reflecting the visual system’s ability to use statistical regularities in the environment to optimize attentional deployment (Theeuwes et al., 2022). Yet it remains unclear whether such preparatory adjustments can extend to modulating the spatial scope of attention based on expectations about upcoming display characteristics.

One domain where preparatory scope adjustment may prove advantageous is visual search, where display density strongly modulates search efficiency. In effortful visual search, response times increase systematically as the number of distractors grows (Treisman & Gelade, 1980; Wolfe, 1994), reflecting the challenge of discriminating targets from distractors when multiple items compete for limited attentional resources. Crucially, dense displays not only increase the number of potential distractor locations but also elevate perceptual demands: targets surrounded by many nearby distractors create crowding effects that impair discrimination (Intriligator & Cavanagh, 2001). Since the zoom lens model posits an inverse relationship between attentional scope and perceptual resolution, concentrating attention in a narrower focus would enhance processing quality within that region, thereby improving target discrimination in crowded conditions. Conversely, in sparse displays with few distractors and minimal crowding, a broader attentional scope might be advantageous, providing coverage across a wider spatial extent without sacrificing discrimination ability. If observers can anticipate whether an upcoming search will be easy or difficult, they might preemptively adjust their attentional zoom lens to optimize performance—narrowing focus when expecting challenging discrimination in dense displays and broadening focus for easier searches.

However, when observers expect distractors to appear in an upcoming search, evidence suggests that preparatory control relies on suppression of distractor locations rather than the signal enhancement mechanisms posited by the zoom lens model. In a behavioral study, Awh et al. (2003) demonstrated that manipulating the probability that visual distractors would appear modulated distractor suppression without concurrent changes in signal enhancement at target locations. Complementing these findings, Serences et al. (2004) used fMRI to show that preparatory attention when expecting high distractor interference selectively enhanced activity in early visual cortex at retinotopic locations corresponding to the expected distractor positions, an effect interpreted as preparatory tagging of distractor locations to facilitate their subsequent suppression. These findings suggest that preparatory control in the context of expected distractors involves location-specific suppression rather than adjustments to the breadth of attentional focus. Nevertheless, these studies employed bilateral cues that directed attention to two locations simultaneously, which may have constrained the ability to observe flexible scope adjustments. Whether observers can proactively narrow or broaden their attentional zoom lens based on expectations about display density—when not constrained by multiple cued locations—thus remains an open question.

To test whether observers flexibly adjust attentional scope based on expected search difficulty, we employed a spatial cueing paradigm in which endogenous cues predicted target location (75% valid) and, critically, we manipulated observers’ expectations about display density across blocks of trials (see Figure 1). At the start of each block, participants were informed that the majority of trials (80%) would contain either sparse displays (1 target, 3 distractors) or dense displays (1 target, 7 distractors). We combined behavioral measures of cueing effects with an inverted encoding model (IEM) applied to EEG data to precisely characterize the spatial selectivity of attention (Foster et al., 2016, 2017). Crucially, this approach allowed us to track not only the locus of attention but also the breadth of attentional resolution with high temporal precision (Feldmann-Wüstefeld & Awh, 2020; van Moorselaar et al., 2025). If observers narrow their attentional zoom lens when expecting difficult search, we should observe sharper spatial tuning profiles in the neural signature of attention during blocks with predominantly dense displays compared to blocks with predominantly sparse displays. This would demonstrate that attentional scope can be proactively adjusted not only in response to cue properties but also based on expected search difficulty, revealing a new dimension of preparatory attentional control.

**Figure 1.**
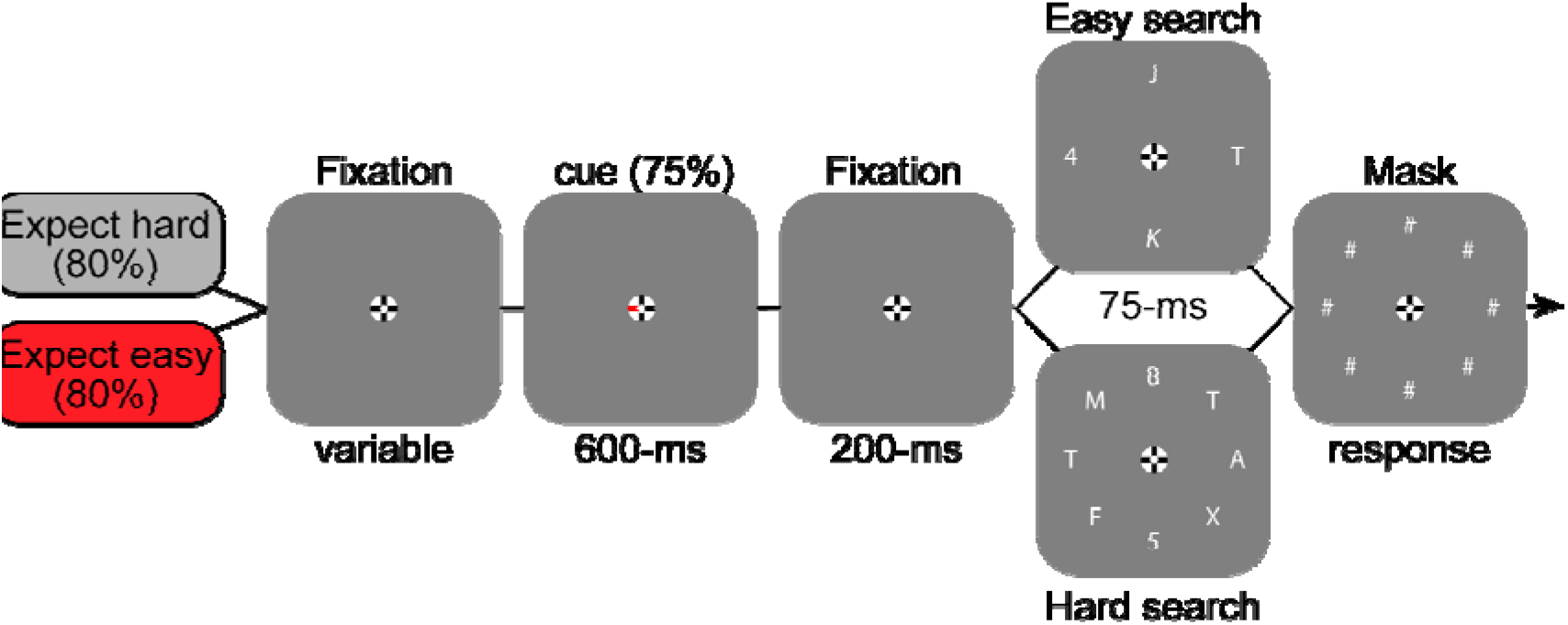
Experimental paradigm. Each trial began with a fixation display followed by a 600-ms spatial cue indicating one of eight possible target locations. After a 200-ms fixation interval, a search display appeared containing either four or eight items (one target digit and distractor letters) for 75 ms, followed by a mask display that remained until response. Masks were presented only at locations that had contained a search item (i.e., four masks in easy displays, eight in hard displays). Participants reported the identity of the target digit via unspeeded keypress. Expectation was manipulated across blocks: in *easy-expectancy* blocks, most trials (80%) contained sparse displays with four items; in *hard-expectancy* blocks, most trials contained dense displays with eight items. The cue predicted the target location with 75% validity.

## Results

### Behavioural results

We examined how expectations about upcoming search difficulty influenced performance in a spatial cueing task. Participants performed visual search across blocks where they expected either sparse (easy) or dense (hard) search displays. On each trial, an endogenous cue directed attention to one of eight possible target locations with 75% validity. Search displays contained either 4 items (easy search) or 8 items (hard search), with the target being a single digit among letter distractors presented briefly (75 ms) before masking (see Figure 1).

As visualized in Figure 2 and revealed by the GLMM (see methods for details) including the predictors Expectation (easy vs hard blocks), Search condition (easy = 4 items vs hard = 8 items) and Cue validity (valid, invalid), performance was better for easy displays displays (β = -1.48, SE = 0.085, z = -17.5, p < .001) and for validly cued trials displays (β = - 1.58, SE = 0.13, z = -11.96, p < .001). The overall main effect of Expectation was not significant (β = 0.12, SE = 0.073, z = 1.64, p = .10). Critically, Expectation interacted with Cue validity (β = -0.25, SE = 0.11, z = -2.36, p = .018), reflecting a larger cueing benefit when dense search displays were expected. To identify the locus of this interaction, we examined the effect of Expectation separately within valid and invalid trials, allowing us to test whether expectations exerted their effects at both cued and uncued locations, or selectively at either cued or uncued locations.

**Figure 2.**
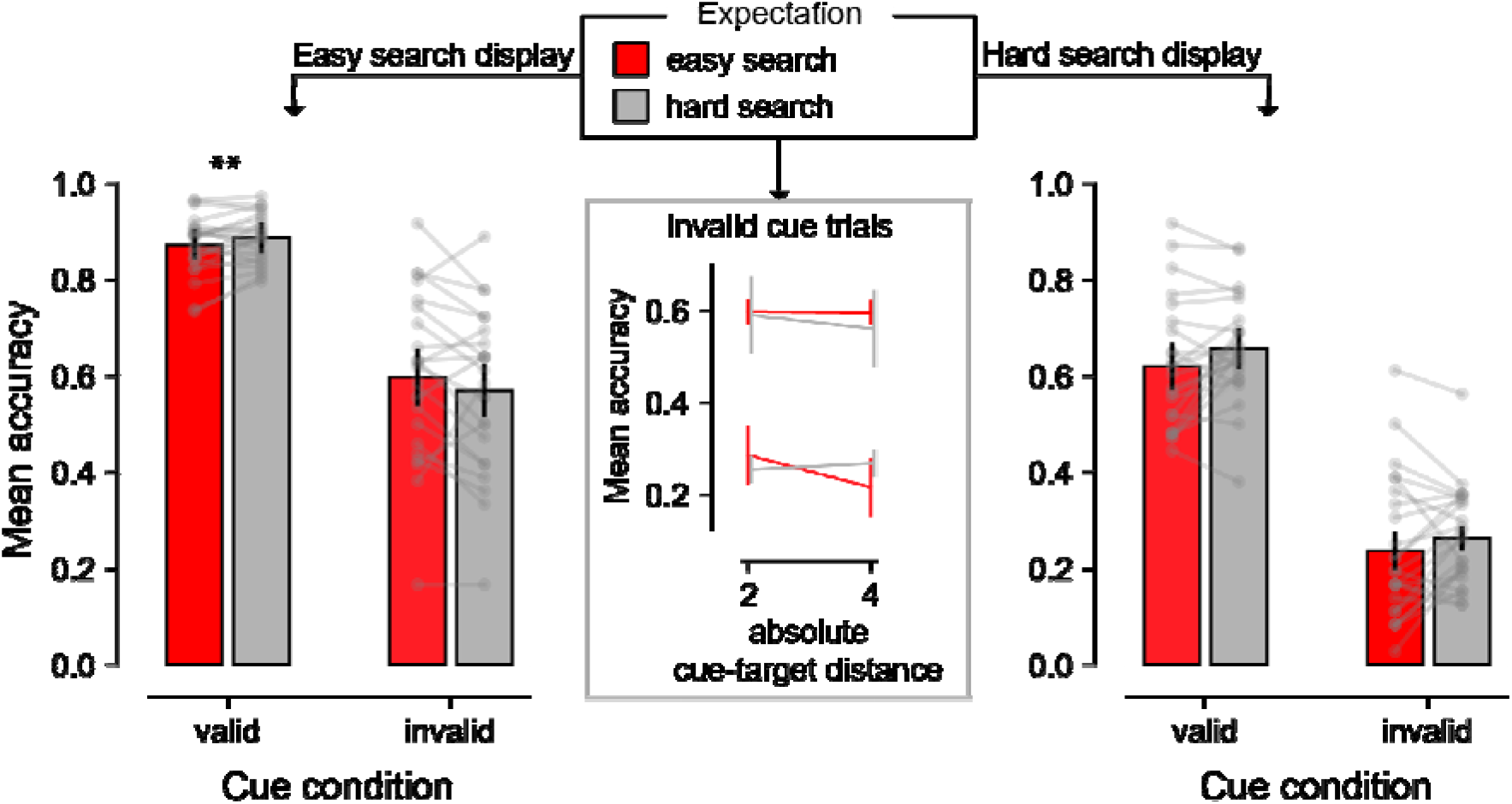
Behavioral results. Mean accuracy for validly and invalidly cued trials as a function of expected search difficulty (easy vs. hard blocks) and search display (easy = 4 items; hard = 8 items). Left and right panels show easy and hard search displays, respectively. Bars represent mean accuracy, with overlaid points indicating individual participant means. Error bars represent 95% within-subject confidence intervals (Morey, 2008). Significance markers reflect planned pairwise comparisons derived from the GLMM using estimated marginal means; only hard search displays showed significantly higher accuracy when participants expected hard (dense) displays (see main text for full statistics). The inset panel shows accuracy on invalid trials as a function of absolute distance between cue and target, with lines corresponding to easy (light) and hard (dark) expectation blocks (see main plot legend); performance declined with increasing distance and was largely unaffected by expectation.

Planned pairwise comparisons using estimated marginal means revealed the locus of this expectation effect. Crucially, on valid trials in hard search displays, where discrimination demands were highest, accuracy was significantly enhanced when participants expected dense displays (odds ratio = 0.86, z = -2.96, *p* = 0.0031). This demonstrates that observers proactively optimized their attentional state for challenging discrimination at the cued location. A similar, though non-significant, pattern emerged for easy displays (odds ratio = 0.89, z = -1.64, *p* = 0.10). By contrast, on invalid trials, Expectation did not significantly affect performance for either display type (easy: odds ratio = 1.14, z = 1.57, *p* = .12; hard: odds ratio = 0.85, z = −1.67, *p* = .094).

A separate GLMM restricted to invalid trials further showed that accuracy decreased with increasing absolute distance from the cued location (Estimate = −0.116, z = −2.98, *p* = .003), with no significant interactions with Search condition or Expectation (all *p*’s > 0.18; see inset Figure 2). Together, these results indicate that Expectation modulated performance primarily at the attended location, while performance at non-cued locations was largely insensitive to expected search difficulty.

### IEM results

To investigate the neural mechanisms underlying the behavioral expectancy effects, we recorded EEG while participants performed the task. We applied an inverted encoding model (IEM) to reconstruct spatially selective CTFs from the scalp distribution of stimulus-evoked broadband EEG activity (Brouwer & Heeger, 2009; Foster et al., 2016). This approach allows us to track how neural populations tuned to different spatial locations respond over time, providing a measure of spatial attention dynamics. Briefly, CTFs characterize the response profile across spatial channels tuned to different locations, and their slope reflects the overall strength of spatially selective neural activity at the attended location, with steeper slopes indicating stronger spatial selectivity (see Methods for full details).

Figure 3A shows that stimulus-evoked CTFs were reliably tuned to the cued location, with tuning emerging around ∼100 ms after cue onset in both expectancy contexts, confirming that the endogenous cues effectively directed spatial attention independent of expectation. As shown in Figure 3B, CTF slope time courses revealed significantly steeper tuning in the *expect-hard* condition than in the *expect-easy* condition (cluster *p* < 0.05), indicating enhanced spatial selectivity when observers anticipated difficult searches.

**Figure 3.**
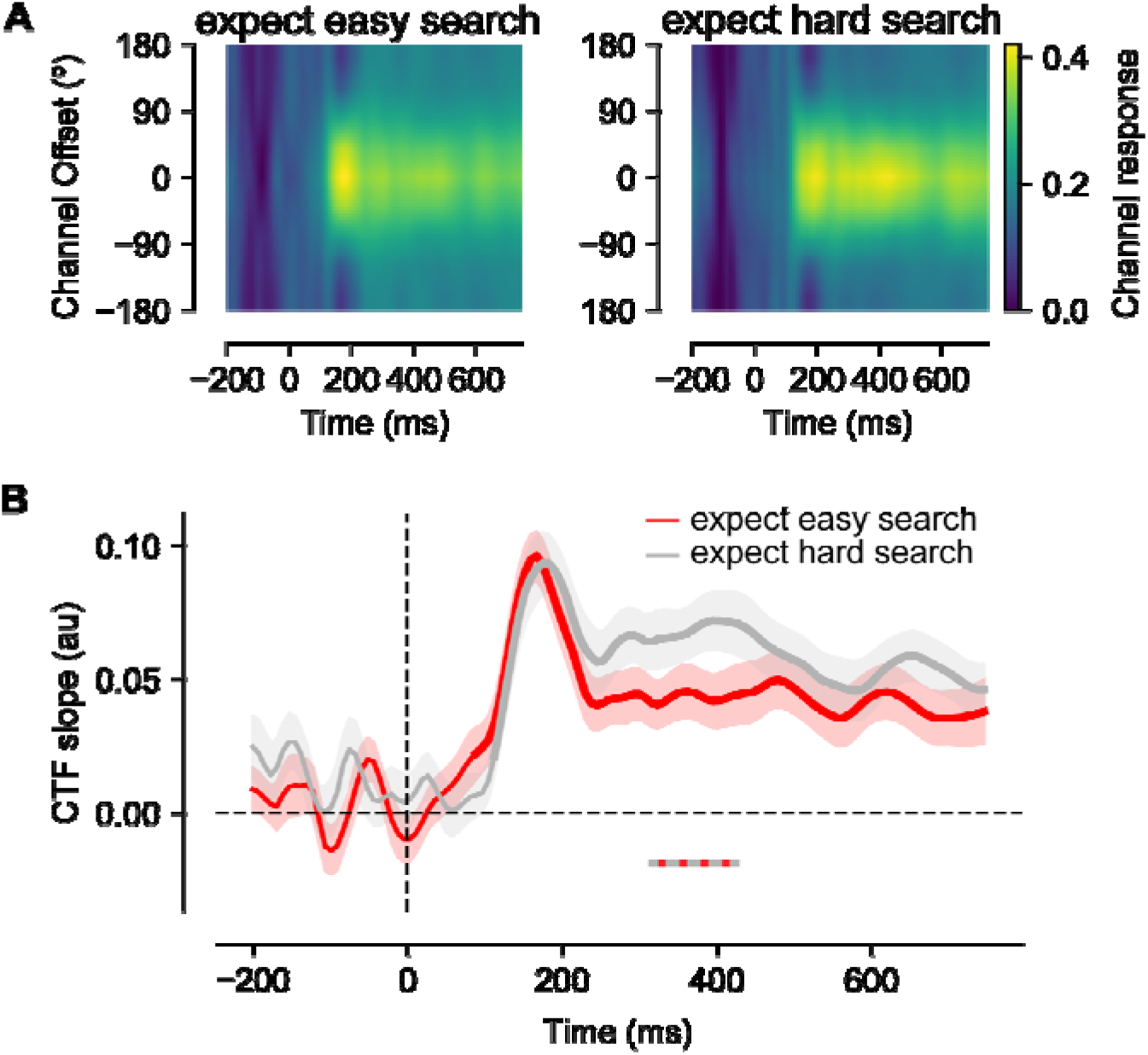
Spatial tuning functions and slope time courses derived from the IEM analysis. **(A)** Reconstructed channel tuning functions (CTFs) for *expect-easy* (left panel) and *expect-hard* (right panel) conditions, averaged across participants. Tuning to the cued location emerged at ∼100 ms following stimulus onset in both conditions, indicating reliable deployment of spatial attention. **(B)** Time course of CTF slopes (mean ± bootsrapped SEM) for each expectancy condition. The dashed grey/red bar along the x-axis marks a significant cluster (cluster-based permutation test, *p* < 0.05) where slopes were steeper in the *expect-hard* condition, indicating enhanced spatial selectivity when participants anticipated difficult search displays.

The steeper CTF slopes in the expect-hard condition appear consistent with the behavioral pattern of enhanced performance at cued locations. However, CTF slopes are agnostic to the underlying neural mechanism: steeper slopes could reflect increased amplitude (higher peak), reduced concentration (narrower tuning), or both. To determine whether, as suggested by the behavioral pattern, this slope difference reflected a gain modulation at the cued location rather than a narrowing of the attentional window, we fitted the CTFs in the significant time window (316–425 ms) with an exponentiated cosine function to separately estimate baseline, amplitude, and concentration (i.e., inverse of tuning width) parameters.

As shown in Figures 4A–B, cue-evoked CTFs within this window exhibited higher amplitude in the *expect-hard* condition (*t* (23) = 2.94, *p* < 0.01), with no corresponding difference in tuning width (*t* (23) = 1.12, *p* = 0.27). To examine the temporal evolution of this amplitude effect, Figure 4C displays the amplitude time course from cue onset through the pre-target interval. Cluster-based permutation testing confirmed that amplitude in the expect-hard search blocks was significantly higher compared to expect-easy search blocks (cluster p < 0.05), while no significant clusters were found for the concentration and baseline parameters. This sustained amplitude enhancement demonstrates that expectations about upcoming search difficulty modulated the *gain* of spatially selective neural activity at the cued location, boosting representational strength in advance of target onset without altering the spatial precision of attentional tuning.

**Figure 4.**
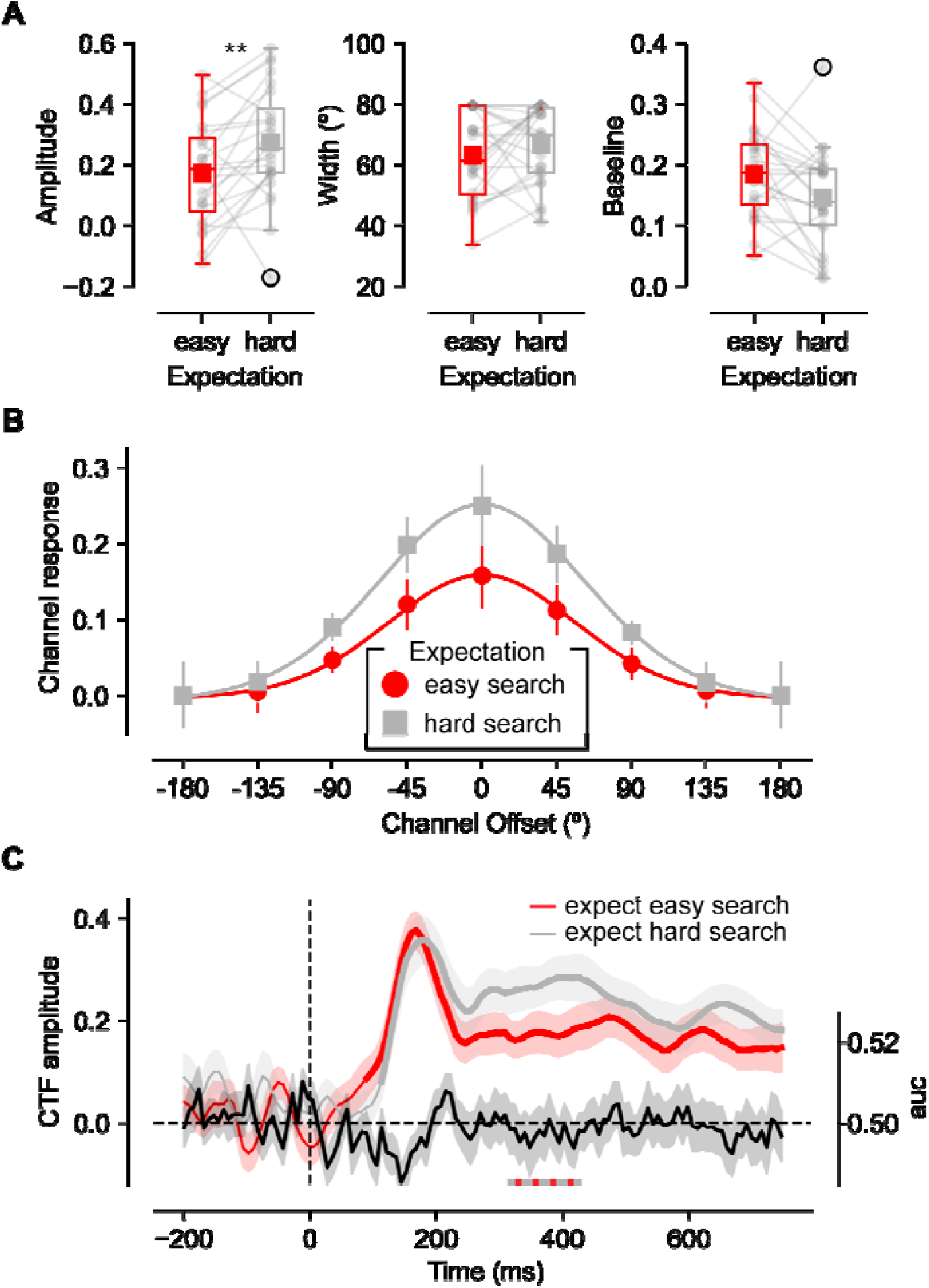
Parameter estimates from CTF fits reveal expectancy-related gain modulation. **(A)** Boxplots showing participant-level estimates of amplitude, concentration (reported as FWHM), and baseline from 316–425 ms after cue onset. Individual data points are overlaid to illustrate between-participant variability; no consistent differences were observed for concentration. Data points beyond 1.5 × IQR are flagged in accordance with the default matplotlib boxplot convention; all participants were retained in the analyses. **(B)** Fitted exponentiated-cosine CTFs from the group-averaged data within the significant time window (316–425 ms), illustrating higher amplitude but comparable width (concentration *k*) in the *expect-hard* condition. Note that baselines were artificially shifted to zero for plotting purposes. **(C)** Time-resolved amplitude estimates (mean ± bootsrapped SEM) from cue onset through the pre-target interval. A significant cluster (*p* < 0.05) indicated higher amplitude in *expect-hard* relative to *expect-easy* blocks, whereas concentration and baseline parameters showed no significant clusters. The black line shows decoding (indexed by auc) of the expected search condition within the same time interval. Together these results indicate that expectancy modulated the gain, but not the spatial precision, of cue-evoked attentional signals.

### Multivariate decoding results

The amplitude enhancement observed in expect-hard blocks raises the question of whether this reflects a quantitative modulation of the same attentional mechanism or the recruitment of a qualitatively different neural process (Koevoet et al., 2025). To address this, we tested whether the expectancy condition (expect-easy vs. expect-hard) could be decoded from the spatial pattern of neural activity following cue onset. If expecting hard versus easy search recruited distinct spatial attention mechanisms, these should manifest as distinguishable spatial patterns that could be classified above chance. However, decoding accuracy remained at chance level throughout the cue-target interval, indicating that the two expectancy conditions did not produce categorically different spatial patterns of neural activity (Figure 4C; black line). It is worth noting that the IEM and decoding analyses address complementary questions: whereas the IEM captures the strength of spatial tuning within each condition, decoding tests whether the spatial pattern of activity differs between conditions. The absence of above-chance decoding is therefore consistent with the IEM findings, as amplitude modulation of a shared spatial pattern would not necessarily produce discriminable multivariate patterns. We note, however, that chance-level decoding should be interpreted with caution, as it may also reflect limited sensitivity of the multivariate approach to detect subtle between-condition differences rather than a true absence of distinct spatial patterns.

These results indicate that expecting difficult versus easy search does not engage qualitatively distinct attentional mechanisms. Instead, both contexts rely on a shared spatially selective process, with expectancy modulating its overall gain rather than its spatial configuration

#### General discussion

The aim of the present study was to examine whether expectations about upcoming search difficulty can proactively modulate the spatial scope of attention. To this end, we combined a spatial cueing paradigm with block-wise manipulations of search expectancy and used an IEM applied to EEG data to reconstruct spatially selective channel tuning functions. Behaviorally, participants showed improved accuracy at the cued location when they expected difficult (dense) search displays, without corresponding costs at uncued locations, suggesting that anticipatory attention selectively benefited the attended region under high expected demands. Consistent with this, at the neural level, expectancy modulated the overall strength of spatially selective activity, with higher CTF amplitudes in the expect-hard condition, while CTF concentration (tuning width) remained unchanged. These findings reveal that expectations about task difficulty enhance the gain of spatial attention rather than narrowing its spatial extent, suggesting that preparatory adjustments operate through amplitude-based gain modulation within an otherwise stable attentional tuning profile.

The present findings extend earlier work on preparatory attentional control by demonstrating that expectations about upcoming search difficulty can modulate attention through proactive signal enhancement. Prior studies manipulating distractor probability (Awh et al., 2003; Serences et al., 2004) concluded that preparatory control predominantly operates via suppression of locations associated with likely distractor occurrence. In those paradigms, attention was divided across two potential target locations, and preparation was defined by whether a distractor was expected to appear or not. Under such circumstances, proactive suppression provides an efficient means of minimizing interference from predictable distractors. In contrast, the present design cued a single target location and varied expectations about overall display density rather than distractor presence. This manipulation emphasized anticipated perceptual demands at the attended site rather than the need to suppress competing locations. Under these conditions, expectation selectively increased activity within the task-relevant spatial channel, consistent with a gain-based enhancement mechanism. Thus, our results suggest that search expectations do not rely exclusively on preparatory suppression but can flexibly operate through enhancement when the task context emphasizes boosting target processing rather than suppressing distractors.

The pattern of enhanced amplitude without concurrent changes in tuning width provides a nuanced perspective on how expectations shape spatial attention. The attentional zoom-lens model has received extensive support for describing how the focus of attention can flexibly narrow or broaden in response to immediate task demands, with a trade-off between spatial extent and processing efficiency (Castiello & Umiltà, 1990; Eriksen & St. James, 1986). However, under the present preparatory conditions, the observed modulation appears not to reflect a change in spatial scope but rather a gain increase within an already established focus of attention. This interpretation is consistent with previous EEG and fMRI work showing that attention can enhance the responsiveness of neurons tuned to the attended location without altering their selectivity profiles (Foster et al., 2021; Itthipuripat, Ester, et al., 2014; Itthipuripat, Garcia, et al., 2014). We note, however, that the absence of a tuning width difference should be interpreted with caution. Subtle or temporally limited changes in attentional scope may not be fully captured by the spatial resolution of the current approach, and we cannot definitively exclude the possibility that expectations also influence spatial extent under some conditions. A related concern is whether the amplitude enhancement observed in expect-hard blocks reflects a spatially specific preparatory mechanism or instead a more general change in alertness or motivational state. Several aspects of the data argue against a purely global account. First, the behavioral benefit of expectation was selective to the cued location, with no corresponding costs or benefits at uncued locations, suggesting that expectancy effects were spatially constrained rather than globally distributed. Second, if expect-hard blocks induced a broadly different neural state through general arousal or motivational engagement, this should manifest as a globally distinct pattern of broadband EEG activity that a multivariate classifier could detect. However, decoding accuracy remained at chance throughout the cue-target interval, indicating that the two expectancy conditions did not produce categorically different spatial patterns of neural activity. Together, these findings suggest that the observed amplitude modulation reflects spatially specific preparatory gain enhancement rather than a global change in task engagement. Thus, while the zoom-lens framework captures flexible adjustments of attentional scope across contexts (Feldmann-Wüstefeld & Awh, 2020; van Moorselaar et al., 2025), our findings suggest that, when preparatory attention is guided by expectations about task difficulty, modulation occurs primarily through amplification of attentional signals rather than through changes in spatial breadth. In this way, gain-based amplification may represent a complementary mechanism by which the visual system optimizes processing efficiency under predictable but demanding conditions.

Our findings extend previous work demonstrating attentional gain at the level of spatial population codes (Foster et al., 2021). Foster and colleagues showed that covert attention increases the amplitude of stimulus-evoked neural responses within ∼100 ms of target onset, reflecting reactive modulation triggered by incoming sensory input. In contrast, we observed a similar gain enhancement proactively, following an endogenous cue pointing to an empty location, with amplitude differences emerging later (∼300 ms). This delay is consistent with the engagement of top-down preparatory mechanisms, as the system interprets the cue and adjusts neural sensitivity in anticipation of upcoming perceptual demands. Together, these findings suggest that the same underlying neural circuitry can implement gain modulation both reactively and proactively, but the timing and trigger of engagement depend on whether attention is stimulus-driven or expectation-driven.

In summary, our results demonstrate that expectations about upcoming search difficulty selectively enhance the strength of spatially selective neural responses at the attended location, without altering their spatial precision. This gain modulation provides a mechanistic explanation for the observed behavioral benefits under high expected difficulty and illustrates that preparatory attention is a flexible, context-sensitive process: the visual system can boost signal strength when needed while maintaining a stable spatial focus.

## Methods

### Participants

The final sample (N = 24; Mean age = ∼22, range = 19 - 28; 16 female; age information of 3 subjects missing), with normal or corrected-to-normal vision, was obtained after replacement of 7 participants, 5 because they failed to hold fixation during the in interstimulus interval (>30% of trials) and 2 because their overall accuracy was below 2.5 S.D. from the group mean. Sample size was based on previous work using inverted encoding models to temporally track covert attention across conditions (Feldmann-Wüstefeld & Awh, 2020; van Moorselaar et al., 2018; van Moorselaar & Slagter, 2019; van Moorselaar et al., 2025). Participants, all with normal or corrected-to-normal vision gave their informed consent prior to the start of the study, which was approved by the University ethical review board. Participants were compensated with course credit or €10 per hour.

### Apparatus, material and procedure

The experiment took place in a dimly lit room on a ASUS ROG STRIX XG248 LED monitor with a 240 Hz refresh rate. Stimuli were presented using Psychopy functionality (Peirce, 2009) within OpenSesame (Mathôt et al., 2012).

Participants were positioned ∼66 cm away from the screen using a desk-mounted chinrest. The eyes were tracked on- and offline using an Eyelink 1000 (SR research) eye tracker tracking the left eye with a 1000 Hz sampling frequency and participants heard a beep each time fixation was broken by more than 1.75° of visual angle in the period before search display onset. At the start of the experiment, eyes were calibrated via a five dots calibration procedure. Drift correction was applied at the start of every new block, and when deemed necessary, the calibration procedure was repeated.

EEG data were recorded at a sampling rate of 512 Hz with default settings using a 32-electrode cap with electrodes placed according to the 10-20 system (Biosemi ActiveTwo system; biosemi.com), with reference electrodes placed at the left and right earlobes. Vertical and horizontal EOG (VEOG/HEOG) were recorded via external electrodes placed ∼2 cm above and below the eye, and ∼1 cm lateral to the external canthi, respectively.

Trials started with a randomly jittered fixation display (1050 – 1350-ms), wherein a fixation marker - a circular shape with an embedded cross hair in black and white (Thaler et al., 2013) - was shown at the center of the screen against a grey background (RGB: 128, 128, 128). This fixation marker was accompanied by eight equidistant, slightly darker grey (RGB: 140, 140, 140) dots (radius = ∼2.7°) in a circular configuration around fixation (radius = ∼0.4°), which remained visible until search display onset. A 600-ms cue display then appeared wherein a red bar within the fixation marker pointed to one of the eight placeholder positions. This cue display was followed by another 200-ms fixation display before search display onset.

The search display consisted of either four or eight equidistant items (height = 25 pixels) placed at the placeholder positions. Each search display contained one randomly selected digit (1–9) and, depending on the search condition, either three or seven letters randomly selected without replacement from the set: A, C, D, E, F, H, K, L, M, N, P, R, S, T, U, V, W, X, Y, Z. The search display was presented for 75 ms before all search stimuli were replaced by pound (#) sign masks. This mask display remained visible until participants indicated via numeric keypress the identity of the target digit. Responses were unspeeded, with emphasis placed on accuracy rather than response time.

Critically, across different experimental blocks, participants expected either sparse or dense displays. In sparse-display blocks, the majority of trials (80%) contained four items (1 target digit + 3 letters) appearing at varying locations across trials to ensure all eight placeholder positions were equally represented. In dense-display blocks, the majority of trials (80%) contained eight items (1 target digit + 7 letters). This expectancy manipulation created two levels of expected search difficulty: sparse displays required relatively easy discrimination with few distractors, while dense displays required more challenging discrimination with many distractors and increased crowding.

In both expectancy conditions, the endogenous cue predicted the target location with 75% probability. On the remaining 25% of trials (i.e., invalid cue trials), the target appeared at one of three distances from the cued location: 2, -2, or 4 steps in the circular space of eight items. Target and cue locations were selected such that they were fully counterbalanced per experimental block.

The experiment started with an 80 trials practice block, out of which 64 trials contained 8 items. Practice continued until accuracy was above 50%. Following practice, participants performed 18 blocks (9 per expectancy condition) of 80 trials each with expectancy switching every three blocks (order counterbalanced). After each block participants received feedback on their performance (i.e., mean accuracy).

### Behavioral analysis

Behavioral accuracy (correct vs. incorrect responses) was analyzed using generalized linear mixed-effects models (GLMMs) with a binomial error distribution and logit link, implemented in R (Team, 2021)(version 4.5.1) using the lme4 package. The Model included fixed effects of Expectation (easy vs. hard blocks), Search condition (easy = 4 items, hard = 8 items), Cue validity (valid vs. invalid), and all interactions. By-participant random intercepts and slopes for all within subject factors were included to account for individual variability. Fixed effects were evaluated using Wald z statistics. Planned comparisons were conducted using estimated marginal means, with contrasts reported as odds ratios and associated z-tests. For continuous predictors (i.e., absolute distance from the cued location on invalid trials), effects are reported as regression coefficients (β), standard errors, z values, and p-values.

### Analysis software

All preprocessing steps and subsequent analytical procedures were executed using custom Python scripts available in a public repository (https://github.com/dvanmoorselaar/DvM). These scripts primarily utilize functionality implemented in the MNE package (Gramfort et al., 2014).

### EEG preprocessing

The recorded EEG data underwent offline re-referencing to the average of both earlobes, followed by zero-phase ‘firwin’ high-pass filtering at 0.01 Hz to eliminate slow signal drifts. Electrodes identified as malfunctioning during recording (*M* = 0.33, range = 0 – 3), were temporarily excluded from analysis to prevent contamination of subsequent preprocessing steps. The continuous recordings were segmented into epochs spanning from 700-ms before to 130-ms after cue display onset, with our primary window of interest being - 200-ms to 800-ms. Following independent component analysis (using MNE’s “picard” method on 1 Hz filtered epochs), eye-blink components were identified and removed.

To eliminate noise-contaminated epochs, we implemented an automated artifact rejection procedure. The EEG signal was band-pass filtered between 110-140 Hz to isolate muscle activity, which was then converted to z-scores. We established participant-specific z-score thresholds based on the variance within the windows of interest. Rather than immediately discarding epochs exceeding this threshold, we employed an iterative approach (Duncan et al., 2023) wherein the five electrodes contributing most substantially to the accumulated z-score within the marked artifact period were identified and sequentially interpolated using spherical splines (Perrin et al., 1989). Only epochs that continued to exceed the threshold after this interpolation procedure were ultimately rejected, resulting in an average exclusion of 4.7% of trials (range: 0.1%-15.8%). In the final preprocessing step, any previously identified malfunctioning electrodes were interpolated using spherical splines

Eye position samples were synchronized with the EEG data during offline analysis and converted to visual degrees representing deviation from central fixation. To maintain interpretative validity, we excluded epochs where gaze position deviated more than 1° from central fixation during the interval from -100 ms to 400 ms relative to cue onset. For instances with missing eye-tracking data, we detected eye movements using an algorithm examining HEOG amplitude changes (window size: 200 ms, step size: 10 ms, threshold: 15 µV). This procedure resulted in the exclusion of an additional 9.5% of the previously cleaned data (range: 2.2%-21.7%).

### Inverted Encoding Model

To reconstruct location-selective channel tuning functions (CTFs) from scalp distributions of frequency power, we implemented an IEM following Foster et al. (2016). This analytical approach assumes that measurements at each electrode reflect the weighted sum of eight spatial channels (neural populations), each selectively tuned to different angular locations. The response profile of each spatial channel was modeled using a half-sinusoid function:

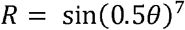

where *θ* represents angular locations (0°-359°) and *R* indicates the channel response. This profile was circularly shifted for each channel to center their peak responses at eight equidistant polar angles (0°, 45°, 90°, etc.), thereby establishing basis functions that specify predicted channel responses across angular locations.

The IEM procedure was applied to 8 Hz low-pass filtered voltages (Bae & Luck, 2019) at each time point through a two-stage process utilizing independent training and test datasets (see *Training and test data below*). In the training stage, we used the training data (*B*_*1*_) to estimate a weight matrix representing each spatial channel’s contribution to electrode-specific power measurements. With *B*_*1*_ (*m* electrodes × *n*_*1*_ observations) representing power values, *C*_*1*_ (*k* channels × *n*_*1*_ observations) denoting predicted channel responses based on basis functions, and *W* (*m* electrodes × *k* channels) characterizing the linear mapping from channel space to electrode space, we formulated a general linear model:

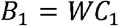

The weight matrix (*W*) was computed using the least squares solution (Python function: np.linalg.lstsq(C_1_, B_1_)). In the testing stage, we inverted the model to transform test data *B*_*2*_ (*m* electrodes × *n*_*2*_ observations) into estimated channel responses *C*_*2*_ (*k* channels × *n*_*2*_ observations) using the derived weight matrix (*Ŵ*) with the Python function np.linalg.lstsq(W.T, B_2_.T).

We circularly shifted these channel responses to center them at a common reference point (0°), corresponding to the cue’s central location. The resulting channel responses (8 channels × 8 location bins) were averaged across location bins to quantify general spatial selectivity in tracking the cued position, independent of specific locations.

### Dataset partitioning

For all IEM analyses, separately for each condition of interest, we randomly divided our data into three independent sets, with two serving as training data (B_1_) and the third as test data (B_2_). Importantly, we used a leave-one-out cross-validation routine, which was repeated until each set had served as testing set (i.e., *B*2). To ensure balance across locations and conditions, we equalized trial numbers across cued locations by selectively discarding excess trials. For each dataset partition, we averaged across trials for individual cue locations to calculate power.

To enhance signal reliability, we implemented an iterative approach for CTF estimation. The procedure of random data partitioning into training and test sets was repeated across 10 iterations, with the resulting CTFs averaged to obtain the final estimates.

### Multivariate decoding

To test whether the two expectancy conditions produced categorically distinct spatial patterns of neural activity, we performed a multivariate pattern analysis using linear discriminant analysis (LDA) with 10-fold cross-validation. The analysis was performed independently at each time sample following cue onset. On each iteration, the classifier was trained to discriminate between expect-easy and expect-hard conditions based on the spatial pattern of activity across all electrodes, with trial counts matched between the two conditions to prevent classification bias. Neural activity was baseline-corrected by subtracting the mean amplitude during the 200 ms window prior to cue onset (-200 to 0 ms). Classification performance was quantified using the area under the receiver operating characteristic curve (AUC), averaged across all cross-validation folds to obtain a single decoding time course for each participant, which was then tested against chance level (0.5) at the group level.

### Statistical Analysis

To evaluate the spatial selectivity of the channel tuning functions (CTFs), we used linear regression to estimate their slopes. Specifically, we calculated the slope of the channel response as a function of location, after collapsing across channels equidistant from the center of the response function.

To better understand how expectation influenced tuning towards the cued location, each CTF was fitted with an exponential cosine function of the form: *f*(*x*) = *e*^*k*(*cos*(*µ*-*x*)-1^)) + *β* , where x is a vector of channel responses, μ, k, and β control the center (i.e., mean), concentration (i.e., inverse of width) and baseline (i.e., vertical offset) of the function, while α corresponds to the amplitude of the function (i.e., vertical scaling) (Ester et al., 2015). These exponential cosine functions were fit separately for expect-easy and expect-hard conditions, yielding separate estimates for the center, baseline, concentration, and amplitude parameters.

For statistical validation of all time course effects, including the reconstructed CTF slopes, fitted CTF parameters, and multivariate decoding performance, we employed non-parametric cluster-based permutation testing. Specifically, we used a one-sample paired t-test in conjunction with Monte Carlo randomization techniques, implemented through MNE’s permutation_cluster_1samp_test function. This statistical approach effectively addresses the temporal correlation inherent in the data while controlling for multiple comparison issues (Maris & Oostenveld, 2007). The statistical procedure followed a resampling protocol wherein each iteration generated a new dataset randomly drawn from the observed data. For each resampled dataset, the signs were randomly flipped, and clusters were formed by identifying adjacent time points with t-values exceeding a predetermined threshold. Only the cluster with the highest cumulative t-value was preserved for each iteration. This resampling and cluster identification process was executed 1024 times (default setting), generating a permutation distribution of cluster statistics. The empirically observed clusters from the actual (non-permuted) data were then evaluated against this permutation distribution. Statistical significance was established when a cluster’s test statistic surpassed the 95^th^ percentile threshold of the permutation distribution, indicating that either the CTF slope itself or the difference between conditions was statistically reliable.

## Acknowledgements

This research was supported by a NWO VICI grant to SVdS. D.v.M designed the study, performed the analyses, and contributed most of the writing. S.S. was closely involved in the design of the experiment, the analysis and made significant contributions to the writing. We would like to thank Tord Helliesen for his invaluable assistance in data collection.

